# Activations of Deep Convolutional Neural Network are Aligned with Gamma Band Activity of Human Visual Cortex

**DOI:** 10.1101/133694

**Authors:** Ilya Kuzovkin, Raul Vicente, Mathilde Petton, Jean-Philippe Lachaux, Monica Baciu, Philippe Kahane, Sylvain Rheims, Juan R. Vidal, Jaan Aru

**Affiliations:** Computational Neuroscience Lab, Institute of Computer Science, University of Tartu, Estonia; INSERM U1028, CNRS UMR5292, Brain Dynamics and Cognition Team, Lyon Neuroscience Research Center, Lyon, France; Université Claude Bernard, Lyon, France; University Grenoble Alpes, LPNC, F-38040 Grenoble, France; CNRS, LPNC UMR 5105, F38040 Grenoble, France; Inserm, U1216, F-38000 Grenoble, France; Neurology Department, CHU de Grenoble, Hôpital Michallon, F-38000 Grenoble, France; Department of Functional Neurology and Epileptology, Hospices Civils de Lyon and Université Lyon, Lyon, France; Epilepsy Institute, Lyon, France; Catholic University of Lyon, France; Department of Penal Law, School of Law, University of Tartu, Estonia

**Author notes:** These authors contributed equally to this work (Raul Vicente, Jaan Aru).

## Abstract

Previous work demonstrated a direct correspondence between the hierarchy of the human visual areas and layers of deep convolutional neural networks (DCNN) trained on visual object recognition. We used DCNNs to investigate which frequency bands correlate with feature transformations of increasing complexity along the ventral visual pathway. By capitalizing on intracranial depth recordings from 100 patients and 11293 electrodes we assessed the alignment between the DCNN and signals at different frequency bands in different time windows. We found that gamma activity, especially in the low gamma-band (30 – 70 Hz), matched the increasing complexity of visual feature representations in the DCNN. These findings show that the activity of the DCNN captures the essential characteristics of biological object recognition not only in space and time, but also in the frequency domain. These results also demonstrate the potential that modern artificial intelligence algorithms have in advancing our understanding of the brain.

**Significance Statement:** Recent advances in the field of artificial intelligence have revealed principles about neural processing, in particular about vision. Previous works have demonstrated a direct correspondence between the hierarchy of human visual areas and layers of deep convolutional neural networks (DCNNs), suggesting that DCNN is a good model of visual object recognition in primate brain. Studying intracranial depth recordings allowed us to extend previous works by assessing when and at which frequency bands the activity of the visual system corresponds to the DCNN. Our key finding is that signals in gamma frequencies along the ventral visual pathway are aligned with the layers of DCNN. Gamma frequencies play a major role in transforming visual input to coherent object representations.

## Introduction

Biological visual object recognition is mediated by a hierarchy of increasingly complex feature representations along the ventral visual stream (DiCarlo et al., 2012). Intriguingly, these transformations are matched by the hierarchy of transformations learned by deep convolutional neural networks (DCNN) trained on natural images (Güçlü and van Gerven, 2015). It has been shown that DCNN provides the best model out of a wide range of neuroscientific and computer vision models for the neural representation of visual images in high-level visual cortex of monkeys (Yamins et al., 2014) and humans (Khaligh-Razavi and Kriegeskorte, 2014). Other studies have demonstrated with fMRI a direct correspondence between the hierarchy of the human visual areas and layers of the DCNN (Güçlü and van Gerven, 2015; Eickenberg et al., 2016; Seibert et al., 2016; Cichy et al., 2016b). In sum, the increasing feature complexity of the DCNN corresponds to the increasing feature complexity occurring in visual object recognition in the primate brain (Kriegeskorte, 2015; Yamins and DiCarlo, 2016).

However, fMRI based studies only allow one to localize object recognition in space, but neural processes also unfold in time and have characteristic spectral fingerprints (i.e. frequencies). With time-resolved magnetoencephalography (MEG) recordings it has been demonstrated that the correspondence between the DCNN and neural signals peaks in the first 200 ms (Cichy et al., 2016b; Seeliger et al., 2017). Here we test the remaining dimension: that biological visual object recognition is also specific to certain frequencies. In particular, there is a long-standing hypothesis that especially gamma band (30 – 150 Hz) signals are crucial for object recognition (Singer and Gray, 1995; Singer, 1999; Fisch et al., 2009; Tallon-Baudry et al., 1997; Tallon-Baudry and Bertrand, 1999; Lachaux et al., 1999; Wyart and Tallon-Baudry, 2008; Lachaux et al., 2005; Vidal et al., 2006; Herrmann et al., 2004; Srinivasan et al., 1999; Levy et al., 2015). More modern views on gamma activity emphasize the role of the gamma rhythm in establishing a communication channel between areas (Fries, 2005, 2015). Further research has demonstrated that especially feedforward communication from lower to higher visual areas is carried by the gamma frequencies (Van Kerkoerle et al., 2014; Bastos et al., 2015; Michalareas et al., 2016). As the DCNN is a feedforward network one could expect that the DCNN will correspond best with the gamma band activity. In this work we used the DCNN as a computational model to assess whether signals in the gamma frequency are more relevant for object recognition than other frequencies.

To empirically evaluate whether gamma frequency has a specific role in visual object recognition we assessed the alignment between the responses of layers of a commonly used DCNN and the neural signals in five distinct frequency bands and three time windows along the areas constituting the ventral visual pathway. Based on the previous findings we expected that: 1) mainly gamma frequencies should be aligned with the layers of the DCNN; 2) the correspondence between the DCNN and gamma should be confined to early time windows; 3) the correspondence between gamma and the DCNN layers should be restricted to visual areas. In order to test these predictions we capitalized on direct intracranial depth recordings from 100 patients with epilepsy and a total of 11293 electrodes implanted throughout the cerebral cortex.

We observed that activity in the gamma range along the ventral pathway is statistically significantly aligned with the activity along the layers of DCNN: gamma (31 – 150 Hz) activity in the early visual areas correlates with the activity of early layers of DCNN, while the gamma activity of higher visual areas is better captured by the higher layers of the DCNN. We also found that while the neural activity in the theta range (5 – 8 Hz) was not aligned with the DCNN hierarchy, the representational geometry of theta activity was correlated with the representational geometry of higher layers of DCNN.

## Materials and Methods

Our methodology involves four major steps described in the following subsections. In “Patients and Recordings” we describe the visual recognition task and data collection. In “Processing of Neural Data” we describe the artifact rejection, extraction of spectral features and the electrode selection processes. “Processing of DCNN Data” shows how we extract activations of artificial neurons of DCNN that occur in response to the same images as were shown to human subjects. In the final step we map neural activity to the layers of DCNN using representational similarity analysis. See Figure 1 for the illustration of the analysis workflow.

**Figure 1.**
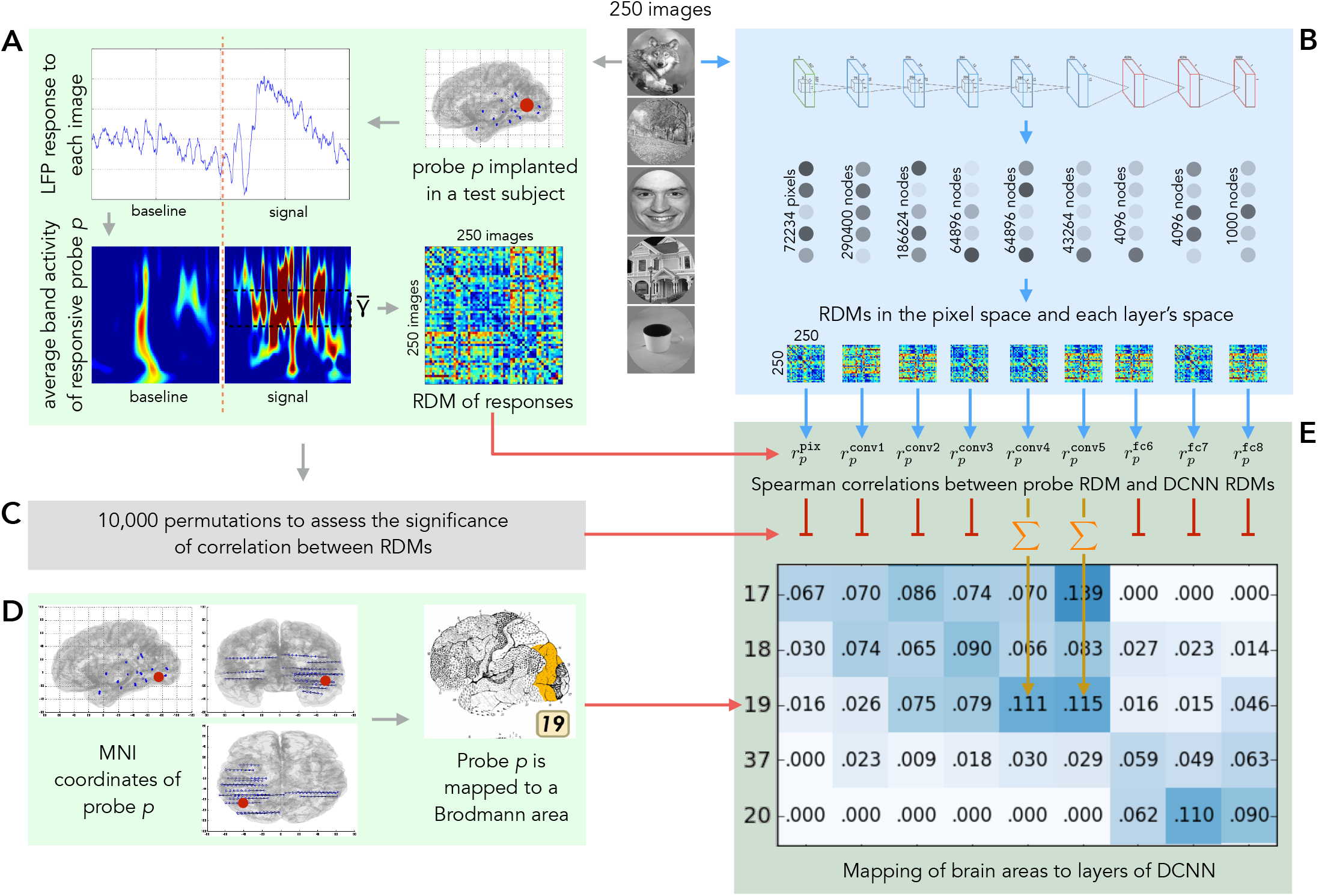
Overview of the analysis pipeline. 250 natural images are presented to human subjects (panel A) and to an artificial vision system (panel B). The activities elicited in these two systems are compared in order to map regions of human visual cortex to layers of deep convolutional neural networks (DCNNs). **A**: LFP response of each of 11293 electrodes to each of the images is converted into the frequency domain. Activity evoked by each image is compared to the activity evoked by every other image and results of this comparison are presented as a representational dissimilarity matrix (RDM). **B**: Each of the images is shown to a pre-trained DCNN and activations of each of the layers are extracted. Each layer’s activations form a representation space, in which stimuli (images) can be compared to each other. Results of this comparison are summarized as a RDM for each DCNN layer. **C**: Subject’s intracranial responses to stimuli are randomly reshuffled and the analysis depicted in panel A is repeated 10000 times to obtain 10000 random RDMs for each electrode. **D**: Each electrode’s MNI coordinates are used to map the electrode to a Brodmann area. The figure also gives an example of electrode implantation locations in one of the subjects (blue circles are the electrodes). **E**: Spearman’s rank correlation is computed between the true (non-permuted) RDM of neural responses and RDMs of each layer of DCNN. Also 10000 scores are computed with the random RDM for each electrode-layer pair to assess the significance of the true correlation score. If the score obtained with the true RDM is significant (the value of *p* < 0.001 is estimated by selecting a threshold such that none of the probes would pass it on the permuted data), then the score is added to the mapping matrix. The procedure is repeated for each electrode and the correlation scores are summed and normalized by the number of electrodes per Brodmann area. The resulting mapping matrix shows the alignment between the consecutive areas of the ventral stream and layers of DCNN.

### Patients and Recordings

100 patients of either gender with drug-resistant partial epilepsy and candidates for surgery were considered in this study and recruited from Neurological Hospitals in Grenoble and Lyon (France). All patients were stereotactically implanted with multilead EEG depth electrodes (DIXI Medical, Besançon, France). The data were bandpass-filtered online from 0.1 to 200 Hz and sampled at 1024 Hz. All participants provided written informed consent, and the experimental procedures were approved by local ethical committee of Grenoble hospital (CPP Sud-Est V 09-CHU-12). Recording sites were selected solely according to clinical indications, with no reference to the current experiment. None of the neurosurgeons who did the operations is among the authors. The authors had no effect on the electrode implantation. The recordings started in 2009, before the present analysis was conceived. All patients had normal or corrected to normal vision.

#### Electrode Implantation

Eleven to 15 semi-rigid electrodes were implanted per patient. Each electrode had a diameter of 0.8 mm and was comprised of 10 or 15 contacts of 2 mm length, depending on the target region, 1.5 mm apart. The coordinates of each electrode contact with their stereotactic scheme were used to anatomically localize the contacts using the proportional atlas of Talairach and Tournoux (Talairach and Tournoux, 1993), after a linear scale adjustment to correct size differences between the patients brain and the Talairach model. These locations were further confirmed by overlaying a post-implantation CT scan (showing contact sites) with a pre-implantation structural MRI with VOXIM^®^ (IVS Solutions, Chemnitz, Germany), allowing direct visualization of contact sites relative to brain anatomy.

All patients voluntarily participated in a series of short experiments to identify local functional responses at the recorded sites (Vidal et al., 2010). The results presented here were obtained from a test exploring visual recognition. All data were recorded using approximately 120 implanted depth electrode contacts per patient with the sampling rates of 512 Hz, 1024 Hz or 2048 Hz. For the current analysis all recordings were downsampled to 512 Hz. Data were obtained in a total of 11293 recording sites.

#### Stimuli and Task

The visual recognition task lasted for about 15 minutes. Patients were instructed to press a button each time a picture of a fruit appeared on screen (visual oddball paradigm). Non-target stimuli consisted of pictures of objects of eight possible categories: houses, faces, animals, scenes, tools, pseudo words, consonant strings, and scrambled images. The target stimuli and last three categories were not included in this analysis. All the included stimuli had the same average luminance. All categories were presented within an oval aperture (illustrated on Figure 1). Stimuli were presented for a duration of 200 ms every 1000 – 1200 ms in series of 5 pictures interleaved by 3-s pause periods during which patients could freely blink. Patients reported the detection of a target through a right-hand button press and were given feedback of their performance after each report. A 2-s delay was placed after each button press before presenting the follow-up stimulus in order to avoid mixing signals related to motor action with signals from stimulus presentation. Altogether, we measured responses to 250 natural images. Each image was presented only once. The images were 3.5 × 4.7 cm on the screen, with a viewing distance of 60-80 cm.

### Processing of Neural Data

The final dataset consists of 2823250 local field potential (LFP) recordings – 11293 electrode responses to 250 stimuli.

To remove the artifacts the signals were linearly detrended and the recordings that contained values ≥ 10*σ_images_*, where *σ_images_* is the standard deviation of responses (in the time window from −500ms to 1000ms) of that particular probe over all stimuli, were excluded from data. All electrodes were re-referenced to a bipolar reference. For every electrode the reference was the next electrode on the same rod following the inward direction. The electrode on the deepest end of each rod was excluded from the analysis. The signal was segmented in the range from −500 ms to 1000 ms, where 0 marks the moment when the stimulus was shown. The −500 to −100 ms time window served as the baseline. There were three time windows in which the responses were measured: 50 − 250 ms, 150 − 350 ms and 250 − 450 ms.

We analyzed five distinct frequency bands: *θ* (5 − 8 Hz), *α* (9 − 14 Hz), *β* (15 − 30 Hz), *γ* (31 − 70 Hz) and Γ (71 − 150 Hz). To quantify signal power modulations across time and frequency we used standard time-frequency (TF) wavelet decomposition (Daubechies, 1990). The signal *s*(*t*) is convoluted with a complex Morlet wavelet *w*(*t*, *f*_0_), which has Gaussian shape in time (*σ_t_*) and frequency (*σ_f_*) around a central frequency fo and defined by *σ_f_* = 1/2*πσ_t_* and a normalization factor. In order to achieve good time and frequency resolution over all frequencies we slowly increased the number of wavelet cycles with frequency (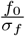 was set to 6 for high and low gamma, 5 for beta, 4 for alpha and 3 for theta). This method allows obtaining better frequency resolution than by applying a constant cycle length (Delorme and Makeig, 2004). The square norm of the convolution results in a time-varying representation of spectral power, given by: *P*(*t, f*_0_) = |*w*(*t, f*_0_)*s*(*t*)|^2^.

Further analysis was done on the electrodes that were responsive to the visual task. We assessed neural responsiveness of an electrode separately for each region of interest – for each frequency band and time window we compared the average post-stimulus band power to the average baseline power with a Wilcoxon signed-rank test for matched-pairs. All p-values from this test were corrected for multiple comparisons across all electrodes with the false discovery rate (FDR) procedure (Genovese et al., 2002). In the current study we deliberately kept only positively responsive electrodes, leaving the electrodes where the post-stimulus band power was significantly weaker than the average baseline power for future work. Table 1 contains the numbers of electrodes that were used in the final analysis in each of 15 regions of interest across the time and frequency domains.

**Table 1.** Number of positively responsive electrodes in each of the 15 regions of interest in a time-resolved spectrogram.

| | $\theta$ | $\alpha$ | $\beta$ | $\gamma$ | $\Gamma$ |
| --- | --- | --- | --- | --- | --- |
| 50 – 250 ms | 1299 | 709 | 269 | 348 | 504 |
| 150 – 350 ms | 1689 | 783 | 260 | 515 | 745 |
| 250 – 450 ms | 1687 | 802 | 304 | 555 | 775 |

Each electrode’s Montreal Neurological Institute coordinate system (MNI) coordinates were mapped to a corresponding Brodmann brain area (Brodmann, 1909) using Brodmann area atlas contained in MRICron (Rorden, 2007) software.

To summarize, once the neural signal processing pipeline is complete, each electrode’s response to each of the stimuli is represented by one number – the average band power in a given time window normalized by the baseline. The process is repeated independently for each time-frequency region of interest.

### Processing of DCNN Data

We feed the same images that were shown to the test subjects to a deep convolutional neural network (DCNN) and obtain activations of artificial neurons (nodes) of that network. We use `Caffe` (Jia et al., 2014) implementation of `AlexNet` (Krizhevsky et al., 2012) architecture (see Figure 8) trained on `ImageNet` (Rus-sakovsky et al., 2015) dataset to categorize images into 1000 classes. Although the image categories used in our experiment are not exactly the same as the ones in the `ImageNet` dataset, they are a close match and DCNN is successful in labelling them.

The architecture of the `AlexNet` artificial network can be seen on Figure 8. It consists of 9 layers. The first is the input layer, where one neuron corresponds to one pixel of an image and activation of that neuron on a scale from 0 to 1 reflects the color of that pixel: if a pixel is black, the corresponding node in the network is not activated at all (value is 0), while a white pixel causes the node to be maximally activated (value 1). After the input layer the network has 5 *convolutional layers* referred to as `convl-5`. A convolutional layer is a collection of filters that are applied to an image. Each filter is a 2D arrangement of weights that represent a particular visual pattern. A filter is convolved with the input from the previous layer to produce the activations that form the next layer. For an example of a visual pattern that a filter of each layer is responsive to, please see the right sub-image under each layer on panel B of Figure 8. Each layer consists of multiple filters and we visualize only one per layer for illustrative purposes. A filter is applied to every possible position on an input image and if the underlying patch of an image coincides with the pattern that the filter represents, the filter becomes activated and translates this activation to the artificial neuron in the next layer. That way, nodes of `conv1` tell us where on the input image each particular visual pattern occurred. Left lower sub-image under each layer on panel B of Figure 8 shows an example output feature map produced by a filter being applied to the input image. Hierarchical structure of convolutional layers gives rise to the phenomenon we are investigating in this work – increase of complexity of visual representations in each sub-sequent layer of the visual hierarchy: in both the biological and artificial systems. Convolutional layers are followed by 3 *fully-connected* layers (`fc6-8`). Each node in a fully-connected layer is, as the name suggests, connected to every node of the previous layer allowing the network to decide which of those connections are to be preserved and which are to be ignored. For both convolutional and fully-connected layers we can apply *deconvolution* (Zeiler and Fergus, 2014) technique to map activations of neurons in those layers back to the input space. This visualization gives better understanding of inner workings of a neural network. Examples of deconvolution reconstruction for each layer are given in the top row of panel B on Figure 8.

For each of the images we store the activations of all nodes of DCNN. As the network has 9 layers we obtain 9 representations of each image: the image itself (referred to as layer 0) in the pixel space and the activation values of each of the layers of DCNN. See panel B of Figure 1 for the cardinalities of those feature spaces.

### Mapping Neural Activity to Layers of DCNN

Once we extracted the features from both neural and DCNN responses our next goal was to compare the two and use a similarity score to map the brain area where a probe was located to a layer of DCNN. By doing that for every probe in the dataset we obtained cross-subject alignment between visual areas of human brain and layers of DCNN. There are multiple deep neural network architectures trained to classify natural images. Our choice of `AlexNet` does not imply that this particular architecture corresponds best to the hierarchy of visual layers of human brain. It does, however, provide a comparison for hierarchical structure of human visual system and was selected among other architectures due to its relatively small size and thus easier interpretability.

Recent studies comparing the responses of visual cortex with the activity of DCNN have used two types of mapping methods. The first type is based on linear regression models that predict neural responses from DCNN activations (Yamins et al., 2014; Güçlü and van Gerven, 2015). The second is based on representational similarity analysis (RSA) (Kriegeskorte et al., 2008). We used RSA to compare distances between stimuli in the neural response space and in the DCNN activation space (Cichy et al., 2016a).

#### Representational Dissimilarity Matrices

We built a representation dissimilarity matrix (RDM) of size *number of stimuli × number of stimuli* (in our case 250 × 250) for each of the probes and each of the layers of DCNN. Note that this is a non-standard approach: usually the RDM is computed over a population (of voxels, for example), while we do it for each probe separately. We use the non-standard approach because often we only had 1 electrode per patient per brain area. Given a matrix RDM^feature space^ a value 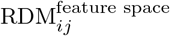 in the *i*th row and *j*th column of the matrix shows the Euclidean distance between the vectors **v**_*i*_ and **v**_*j*_ that represent images *i* and *j* respectively in that particular feature space. Note that the preprocessed neural response to an image in a given frequency band and time window is a scalar, and hence correlation distance is not applicable. Also, given that DCNNs are not invariant to the scaling of the activations or weights in any of its layers, we preferred to use closeness in Euclidean distance as a more strict measure of similarity. In our case there are 10 different feature spaces in which an image can be represented: the original pixel space, 8 feature spaces for each of the layers of the DCNN and one space where an image is represented by the preprocessed neural response of probe *p*. For example, to analyze region of interest of high gamma in 50 – 250 ms time window we computed 504 RDM matrices on the neural responses – one for each positively responsive electrode in that region of interest (see Table 1), and 9 RDM matrices on the activations of the layers of DCNN. A pair frequency band and a time window, such as “high gamma in 50-250 ms window” is referred to as *region of interest* in this work.

#### Representational Similarity Analysis

The second step was to compare the RDM^probe *p*^ of each probe *p* to RDMs of layers of DCNN. We used Spearman’s rank correlation as measure of similarity between the matrices:

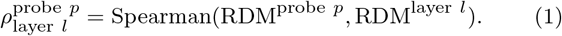

As a result of comparing RDM^probe *p*^ with every RDM^layer *l*^ we obtain a vector with 9 scores: (*ρ*_pixels_, *ρ*_conv1_,…,*ρ*_fc8_) that serves as a distributed mapping of probe p to the layers of DCNN (see panel E of Figure 1). The procedure is repeated independently for each probe in each region of interest. To obtain an aggregate score of the correlation between an area and a layer the *ρ* scores of all individual probes from that area are summed and divided by the number of *ρ* values that have passed the significance criterion. The data for the Figures 3 and 5 are obtained in such manner.

Figure 2 presents the results of applying RSA within the DCNN to compare the similarity of representational geometry between the layers.

**Figure 2.**
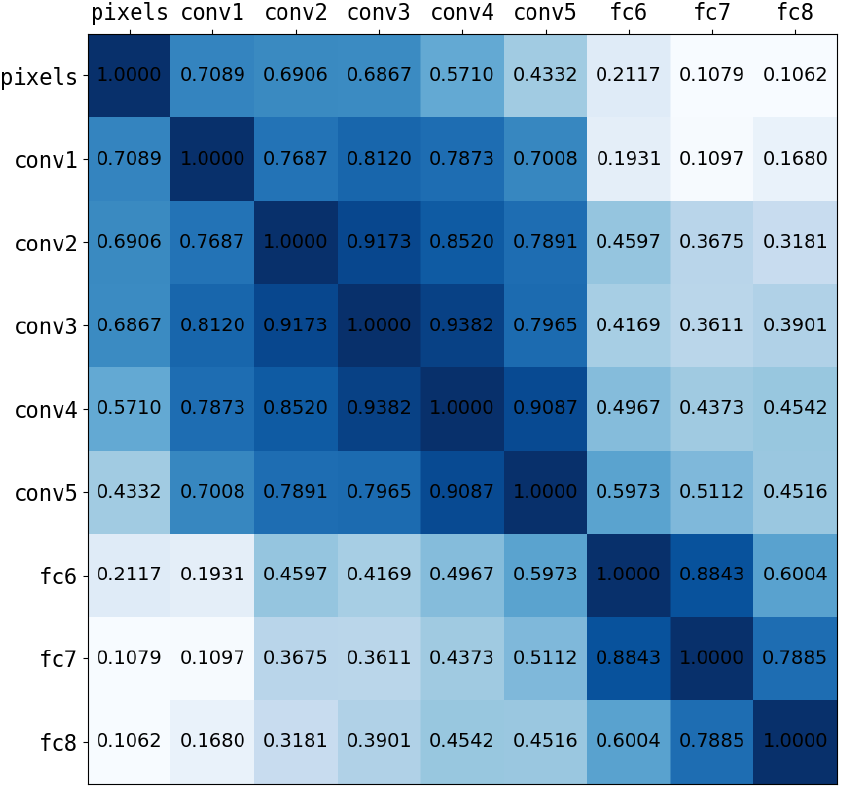
Correlations between the RDMs of DCNN layers. All scores are significant.

#### Statistical significance and controls

To assess the statistical significance of the correlations between the RDM matrices we ran a permutation test. In particular, we reshuffled the vector of brain responses to images 10000 times, each time obtaining a dataset where the causal relation between the stimulus and the response is destroyed. On each of those datasets we ran the analysis and obtained Spearman’s rank correlation scores. To determine score’s significance we compared the score obtained on the original (unshuffled) data with the distribution of scores obtained with the surrogate data. If the score obtained on the original data was bigger than the score obtained on the surrogate sets with *p* < 0.001 significance, we considered the score to be significantly different. The threshold of *p* = 0.001 is estimated by selecting such a threshold that on the surrogate data none of the probes would pass it.

To size the effect caused by training artificial neural network on natural images we performed a control where the whole analysis pipeline depicted on Figure 1 is repeated using activations of a network that was not trained – its weights are randomly sampled from a Gaussian distribution 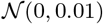.

For the relative comparison of alignments between the bands and the noise level estimation we took 1,000 random subsets of half of the size of the dataset. Each region of interest was analyzed separately. The alignment score was calculated for each subset, resulting in 1,000 alignment estimates per region of interest. This allowed us to run a statistical test between each pair of regions of interest to test the hypothesis that the DCNN alignment with the probe responses in one band is higher than the alignment with the responses in another band. We used Mann-Whitney U test (Mann and Whitney, 1947) to test that hypothesis and accepted the difference as significant at p-value threshold of 0.005 Bonferroni corrected (Dunn, 1961) to 2.22e−5.

### Quantifying properties of the mapping

To evaluate the results quantitatively we devised a set of measures specific to our analysis. *Volume* is the total sum of significant correlations (see Equation 1) between the RDMs of the subset of layers *L* and the RDMs of the probes in the subset of brain areas *A*:

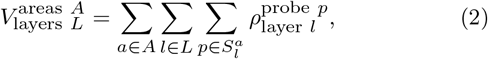

where *A* is a subset of brain areas, *L* is a subset of layers, and 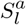 is the set of all probes in area *a* that significantly correlate with layer *l*.

We express *volume of visual activity* as

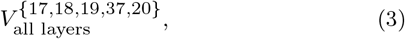

which shows the total sum of correlation scores between all layers of the network and the Brodmann areas that are located in the ventral stream: 17, 18, 19, 37, and 20.

*Visual specificity* of activity is the ratio of volume in visual areas and volume in all areas together, for example visual specificity of all of the activity in the ventral stream that significantly correlates with any of layers of DCNN is

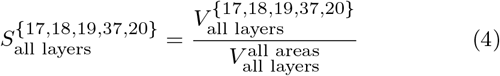

The measures so far did not take into account hierarchy of the ventral stream nor the hierarchy of DCNN. The following two measures are the most important quantifiers we rely on in presenting our results and they do take hierarchical structure into account.

The *ratio of complex visual features to all visual features* is defined as the total volume mapped to layers `conv5, fc6, fc7` divided by the total volume mapped to layers `convl, conv2, conv3, conv5, fc6, fc7`:

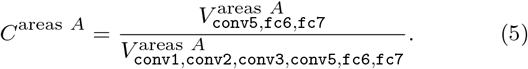

Note that for this measure layers `conv4` and `fc8` are omitted: layer `conv4` is considered to be the transition between the layers with low and high complexity features, while layer `fc8` directly represents class probabilities and does not carry visual representations of the stimuli (if only on very abstract level).

Finally, the *alignment* between the activity in the visual areas and activity in DCNN is estimated as Spearman’s rank correlation between two vectors each of length equal to the number of probes with RDMs that significantly correlate with an RDM of any of DCNN layers. The first vector is a list of Brodmann areas BA^*p*^ where each of such probes *p* belongs:

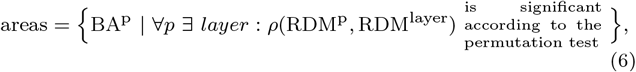

ordered by the hierarchy of the ventral stream: BA17, BA18, BA19, BA37, BA20. Areas are coded by integer range from 0 to 4. The second vector lists DCNN layers L^*p*^ to which the very same probes *p* were assigned:

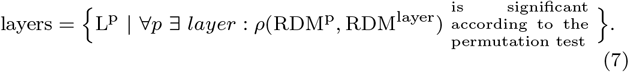

Layers of DCNN are coded by integer range from 0 to 8. We denote Spearman rank correlation of those two vectors as *alignment*

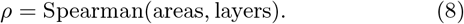

We note that although the hierarchy of the ventral stream is usually not defined through the progression of Brodmann areas, such ordering nevertheless provides a reasonable approximation of the real hierarchy (Lerner et al., 2001; Grill-Spector and Malach, 2004). As both the ventral stream and the hierarchy of layers in DCNN have an increasing complexity of visual representations, the relative ranking within the biological system should coincide with the ranking within the artificial system. Based on the recent suggestion that significance levels should be shifted to 0.005 (Dienes et al., 2017) and after Bonferroni-correcting for 15 time-frequency windows we accepted alignment as significant when it passed *p* < 0.0003(3).

### Data and code availability

The full code of the analysis pipeline is publicly available at https://github.com/kuz/Human-Intracranial-Recordings-and-DCNN-to-Compare-Biological-and-Artificial-Mechanisms-of-Vision. All raw human brain recordings that support the findings of this study are available from Lyon Neuroscience Research Center but restrictions apply to the availability of these data, which were used under license for the current study, and so are not publicly available. Raw data are however available from the authors upon reasonable request and with permission of Lyon Neuroscience Research Center. All the pre-processed data are available and download links are provided in the documentation of the code repository referenced above.

## Results

### Activity in gamma band is aligned with the DCNN

We tested the hypothesis that gamma activity has a specific role in visual object recognition compared to other frequencies. To that end we assessed the alignment of neural activity in different frequency bands and time windows to the activity of layers of a deep convolutional neural network (DCNN) trained for object recognition.

In particular, we used RSA to compare the representational geometry of different DCNN layers and the activity patterns of different frequency bands of single electrodes (see Figure 1). We consistently found that signals in low gamma (31 – 70 Hz) frequencies across all time windows and high gamma (71 – 150 Hz) frequencies in 150 – 350 ms window are aligned with the DCNN in a specific way: increase of the complexity of features along the layers of the DCNN was roughly matched by the transformation in the representational geometry of responses to the stimuli along the ventral stream. In other words, the lower and higher layers of the DCNN explained gamma band signals from earlier and later visual areas, respectively.

Figure 3 illustrates assignment of neural activity in low gamma band (panel A) and high gamma band (panel B) to Brodmann areas and layers of DCNN. As one can see, most of the activity was assigned to visual areas (areas 17, 18, 19, 37, 20). Focusing on visual areas (panels C, D) revealed a diagonal trend that illustrates the alignment between ventral stream and layers of DCNN. Our findings across all subjects, time windows and frequency bands are summarized on the left panel of Figure 4. We note that the alignment in the gamma bands is also present at the single-subject level as can be seen in Figure 6.

**Figure 3.**
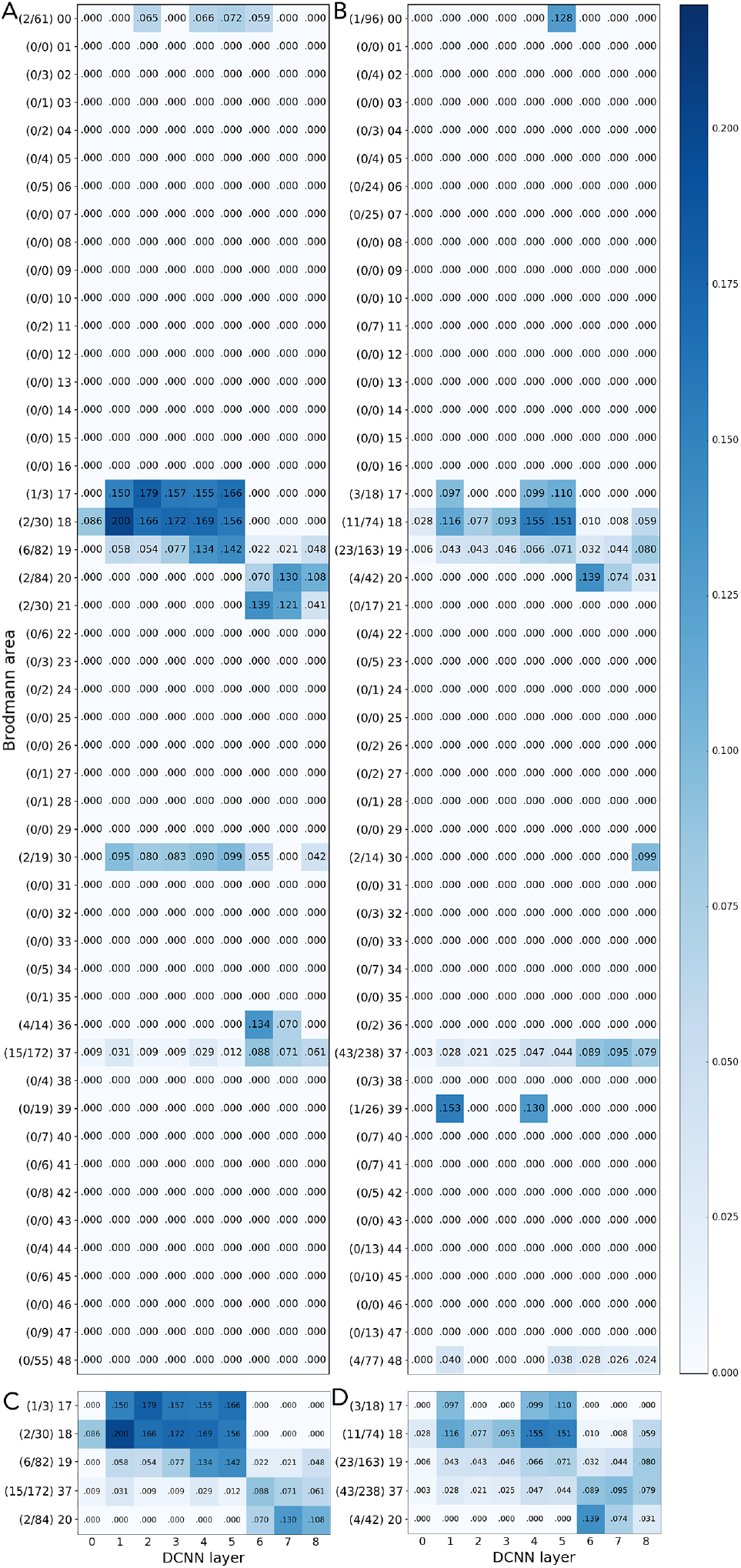
Mapping of the activity in Brodmann areas to DCNN layers. Underlying data comes from the activity in low gamma (31-70 Hz, subfigures A and C) and high gamma (71-150 Hz, subfigures B and D) bands in 150-350 ms time window. C and D are subselection of the areas that constitute ventral stream: 17, 18, 19, 37, 20. On the vertical axis there are Brodmann areas and the number of significantly correlating probes in each area out of the total number of responsive probes in that area. Horizontal axis represents succession of layers of DCNN. Number in each cell of the matrix is the total sum of correlations (between RDMs of probes in that particular area and the RDM of that layer) normalized by the number of significantly correlating probes in an area. There are two important observations to be made out of this plot: a) statistically significant neural responses are specific to visual areas, b) the alignment between the ventral stream and layer of DCNN is clearly visible. Area 0 contains the regions of the brain not mapped by the atlas.

**Figure 4.**
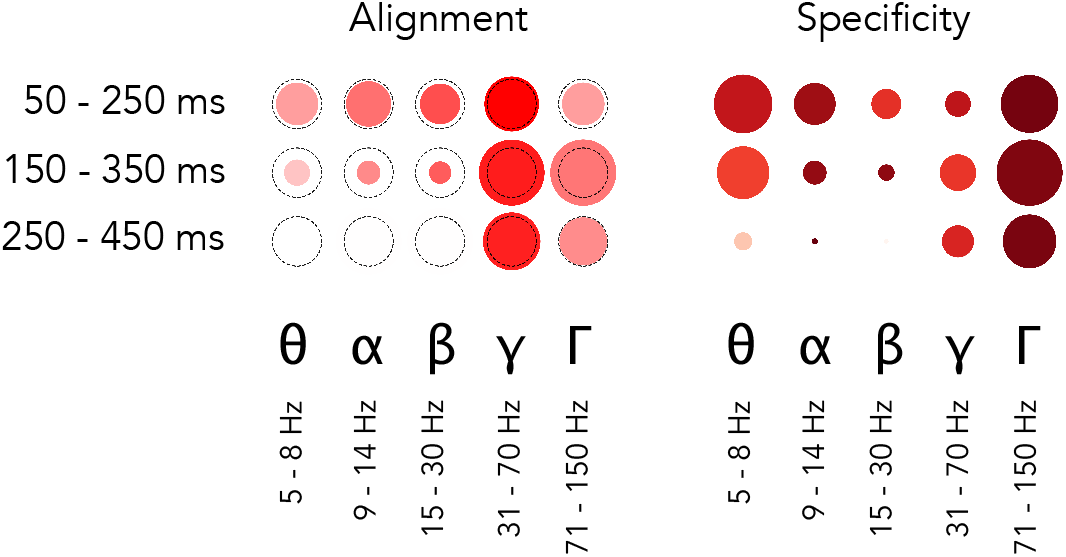
Overall relative statistics of brain responses across frequency bands and time windows. The left panel shows the alignment between visual brain areas and DCNN layers (see Equation 8). The color indicates the correlation value (*ρ*) while the size of the marker shows the logarithm (so that not significant results are still visible on the plot) of inverse of the statistical significance of the correlation, dotted circle indicates *p* = 0.0003(3) – the Bonferroni-corrected significance threshold level of 0.005. The right panel shows whether activity in a region of interest is specific to visual areas (see Equation 4): intense red means that most of the activity in that band and time window happened in visual areas, size of the marker indicates total volume (Equation 2) of activity in all areas. The maximal size of a marker is defined by the biggest marker on the figure.

Apart from the alignment we looked at the total amount of correlation and its specificity to visual areas. On the right panel of Figure 4 we can see that the volume of significantly correlating activity was highest in the high gamma range. Remarkably, 97% of that activity was located in visual areas, which is confirmed by Figure 3 where we see that in the gamma range only a few electrodes were assigned to Brodmann areas that are not part of the ventral stream.

### Activity in other frequency bands

To test the specificity of gamma frequency in visual object recognition, we assessed the alignment between the DCNN and other frequencies. The detailed mapping results for all frequency bands and time windows are presented in layer-to-area fashion on Figure 5. The results on the right part of Table 2 show the alignment values and significance levels for a DCNN that is trained for object recognition on natural images. On the left part of Table 2 the alignment between the brain areas and a DCNN that has not been trained on object recognition (i.e. has random weights) is given for comparison. One can see that training a network to classify natural images drastically increases the alignment score ρ and its significance. One can see that weaker alignment (that does not survive the Bonferroni correction) is present in early time window in theta and alpha frequency range. No alignment is observed in the beta band.

**Figure 5.**
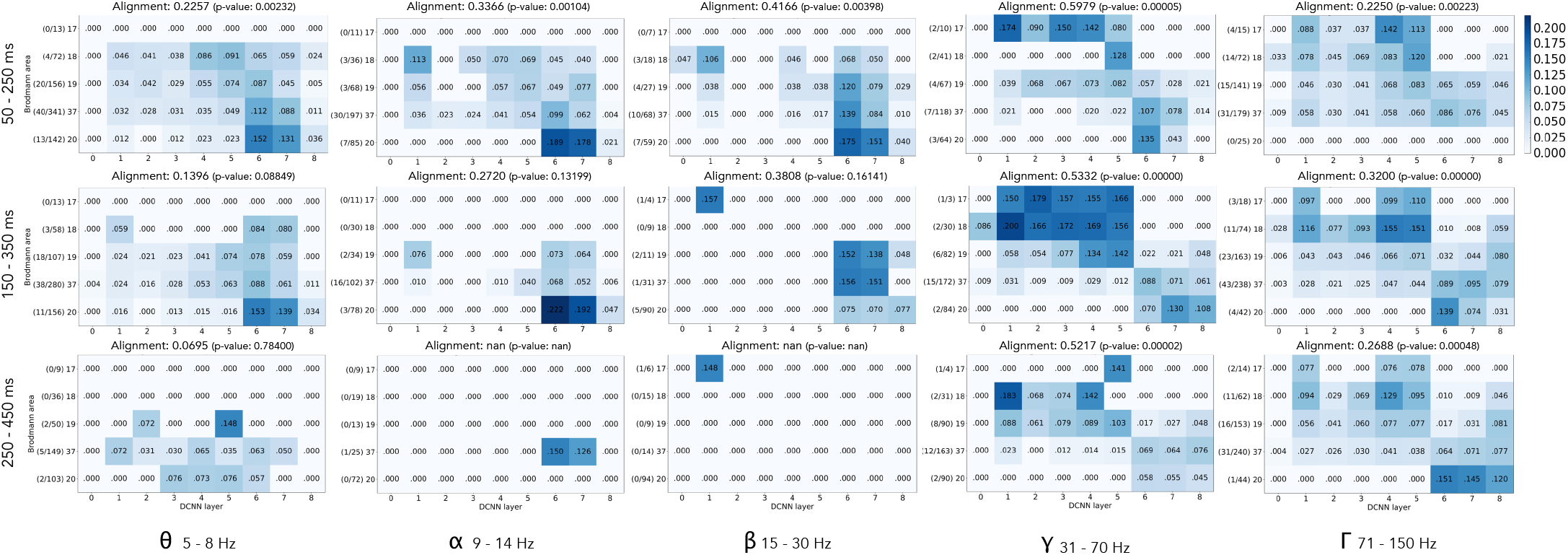
Mapping of activity in visual areas to activations of layers of DCNN across five frequency bands and three time windows. Vertical axis holds Brodmann areas in the order of the ventral stream (top to bottom), horizontal axis represents the succession of layers of DCNN. Number in each cell of a matrix is the total sum of correlations (between RDMs of probes in that particular area and the RDM of that layer) normalized by the number of significantly correlating probes in an area. The alignment score is computed as Spearman’s rank correlation between electrode assignment to Brodmann areas and electrode assignment to DCNN layers (Equation 8). The numbers on the left of each subplot show the number of significantly correlating probes in each area out of the total number of responsive probes in that area.

**Table 2.** Alignment *ρ* score and significance for all 15 regions of interest. * indicates the alignments that pass p-value threshold of 0.05 Bonferroni-corrected to 0.003(3) and *** the ones that pass 0.005 (Dienes et al., 2017) Bonferroni-corrected to 0.0003(3). Note how the values differ between random (control) network and a network trained on natural images. Visual representation of alignment and significance is given on the left panel of Figure 4.

| Band | Window | Alignment with layers of randomly initialized AlexNet |  | Alignment with layers of AlexNet trained on ImageNet |  |  |
| --- | --- | --- | --- | --- | --- | --- |
| | | $\rho$ | p-value | $\rho$ | p-value | |
| $\theta$ | 50-250 ms | 0.0632 | 0.71 | 0.2257 | 0.00231575 | * |
| $\theta$ | 150-350 ms | -0.1013 | 0.59 | 0.1396 | 0.08848501 | |
| $\theta$ | 250-450 ms | 0.1396 | 0.59 | 0.0695 | 0.78400416 | |
| $\alpha$ | 50-250 ms | -0.2411 | 0.32 | 0.3366 | 0.00103551 | * |
| $\alpha$ | 150-350 ms | 0.0000 | 1.00 | 0.2720 | 0.13199463 | |
| $\alpha$ | 250-450 ms | — | — | — | — | |
| $\beta$ | 50-250 ms | — | — | 0.4166 | 0.00397929 | |
| $\beta$ | 150-350 ms | — | — | 0.3808 | 0.16141286 | |
| $\beta$ | 250-450 ms | — | — | — | — | |
| $\gamma$ | 50-250 ms | 0.1594 | 0.62 | 0.5979 | 0.00004623 | *** |
| $\gamma$ | 150-350 ms | -0.1688 | 0.34 | 0.5332 | 0.00000059 | *** |
| $\gamma$ | 250-450 ms | -0.1132 | 0.56 | 0.5217 | 0.00001624 | *** |
| $\Gamma$ | 50-250 ms | 0.0869 | 0.42 | 0.2259 | 0.00222940 | * |
| $\Gamma$ | 150-350 ms | -0.0053 | 0.96 | 0.3200 | 0.00000051 | *** |
| $\Gamma$ | 250-450 ms | -0.1361 | 0.33 | 0.2688 | 0.00047999 | * |

In order to take into account the intrinsic variability when comparing alignments of different bands between each other, we performed a set of tests to see which bands have statistically significantly higher alignment with DCNN than other bands. See the methods section on “Statistical significance and controls” for details. The results of those tests are presented in Table 3. We draw the following conclusions from these results: (a) alignment of DCNN with low gamma (31 – 70 Hz) band is larger than the alignment with any other band, (b) within the low gamma the activity in early time window 50 – 250 ms is aligned more that in later windows, (c) alignment in the high gamma (71 – 150 Hz) is higher that alignment of *θ*, but not higher than alignment of *α*, (d) within the high gamma band the activity in the middle time window 150 – 350 ms has the highest alignment, followed by late 250 – 450 ms window and then by the early activity in 50 – 250 ms window, (e) theta band has the weakest alignment across all bands, (f) alignment of early alpha activity is higher than the alignment of early and late high gamma.

**Table 3.** Alignment of the region of interest on the left is statistically significantly larger than the alignments of the regions of interest on the right. To obtain these results a pairwise comparison of the magnitude of alignment between the regions of interest was made. First column enlists significantly aligned regions, their average alignment *ρ* score when estimated on 1000 random subsets of the data (each of the half of the size of the dataset), and standard deviation of the alignment. On the right side of the table we list the regions of interest of which the ROI on the left is larger. The hypothesis was tested using Mann-Whithney U test and only the results with the p-values that have passed the threshold of 2.2e–5 (0.005 Bonferroni corrected to take into account multiple comparisons) are presented in the table.

|  |  |  |  |
| --- | --- | --- | --- |
| $0.2079 \pm 0.1381$ | $\theta^{50}$ | > | - |
| $0.3352 \pm 0.0989$ | $\alpha^{50}$ | > | $\theta^{50}, \Gamma^{50}, \Gamma^{250}$ |
| $0.5652 \pm 0.1953$ | $\gamma^{50}$ | > | $\theta^{50}, \alpha^{50}, \gamma^{150}, \gamma^{250}, \Gamma^{50}, \Gamma^{150}, \Gamma^{250}$ |
| $0.4880 \pm 0.1650$ | $\gamma^{150}$ | > | $\theta^{50}, \alpha^{50}, \Gamma^{50}, \Gamma^{150}, \Gamma^{250}$ |
| $0.4656 \pm 0.2185$ | $\gamma^{250}$ | > | $\theta^{50}, \alpha^{50}, \Gamma^{50}, \Gamma^{150}, \Gamma^{250}$ |
| $0.2172 \pm 0.1179$ | $\Gamma^{50}$ | > | - |
| $0.3116 \pm 0.1115$ | $\Gamma^{150}$ | > | $\theta^{50}, \Gamma^{50}, \Gamma^{250}$ |
| $0.2494 \pm 0.1381$ | $\Gamma^{250}$ | > | $\theta^{50}, \Gamma^{50}$ |

### The observed alignment with the DCNN is dependent of having two groups of layers in its architecture

On Figures 2 and 4 one can observe that sites in lower visual areas (17,18) are mapped to DCNN layers 1 to 5 without a clear trend but are not mapped to layers 6-8. Similarly areas 37 and 20 are mapped to layers 6-8 but not to 1-5. Hence we next asked whether the observed alignment is depending on having two different groups of visual areas related to two groups of DCNN layers. We tested this by computing alignment within the subgroups. We looked at alignment only between the lower visual areas (17,18,19) and the convolutional layers 1-5, and separately at the alignment between higher visual areas (37, 20) and fully connected layers of DCNN (6-8). We observed no significant alignment within any of the subgroups. So we conclude that the alignment mainly comes from having different groups of areas related more or less equally to two groups of layers. The underlying reason for having these two groups of layers comes from the structure of the DCNN - it has two different types of layers, convolutional (layers 1-5) and fully connected (layers 6-8) (See Figure 8 panels A and B for a visualization of the different layers and their learned features and Discussion section “Two groups of areas in the visual system are mapped to two groups of layers in the DCNN” for a longer explanation of the differences of these layers). As can be evidenced on Figure 2 the layers 1-5 and 6-8 of the DCNN indeed cluster into two groups. Taken together, we observed that early visual areas are mapped to the convolutional layers of the DCNN whereas higher visual areas match the activity profiles of the fully connected layers of the DCNN.

**Figure 6.**
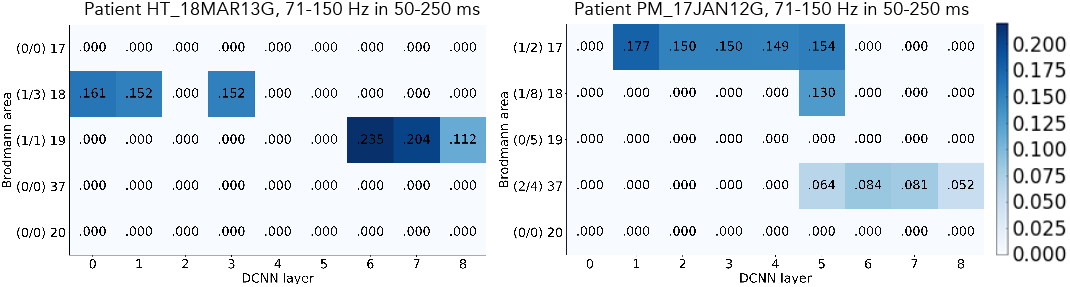
Single subject results from two different subjects. The numbers show the sum of correlations normalized by the number of probes in an area. On the left plot we see how a probe in Brodmann area 18 is mapped to the layers 0, 1, and 3 DCNN, while the activity in Brodmann area 19, which is located further along the ventral stream, is mapped to the higher layers of DCNN: 6, 7, 8. Similar trend is seen on the right plot. The numbers on the left of each subplot show the number of significantly correlating probes in each area out of the total number of responsive probes in that area.

### Complexity of visual features is reflected by different visual areas and frequencies

To investigate the involvement of each frequency band more closely we analyzed each visual area separately. Figure 7 shows the volume of activity in each area (size of the marker on the figure) and whether that activity was more correlated with the complex visual features (red color) or simple features (blue color). In our findings the role of the earliest area (17) was minimal, however that might be explained by a very low number of electrodes in that area in our dataset (less than 1%). One can see on Figure 7 that activity in theta frequency in time windows 50 – 250 ms and 150 – 350 ms had large volume and is correlated with the higher layers of DCNN in higher visual areas (19, 37, 20) of the ventral stream. This hints at the role of activity reflected by the theta band in visual object recognition. In general, in areas 37 and 20 all frequency bands reflected the information about high level features in the early time windows. This implies that already at early stages of processing the information about complex features was present in those areas.

**Figure 7.**
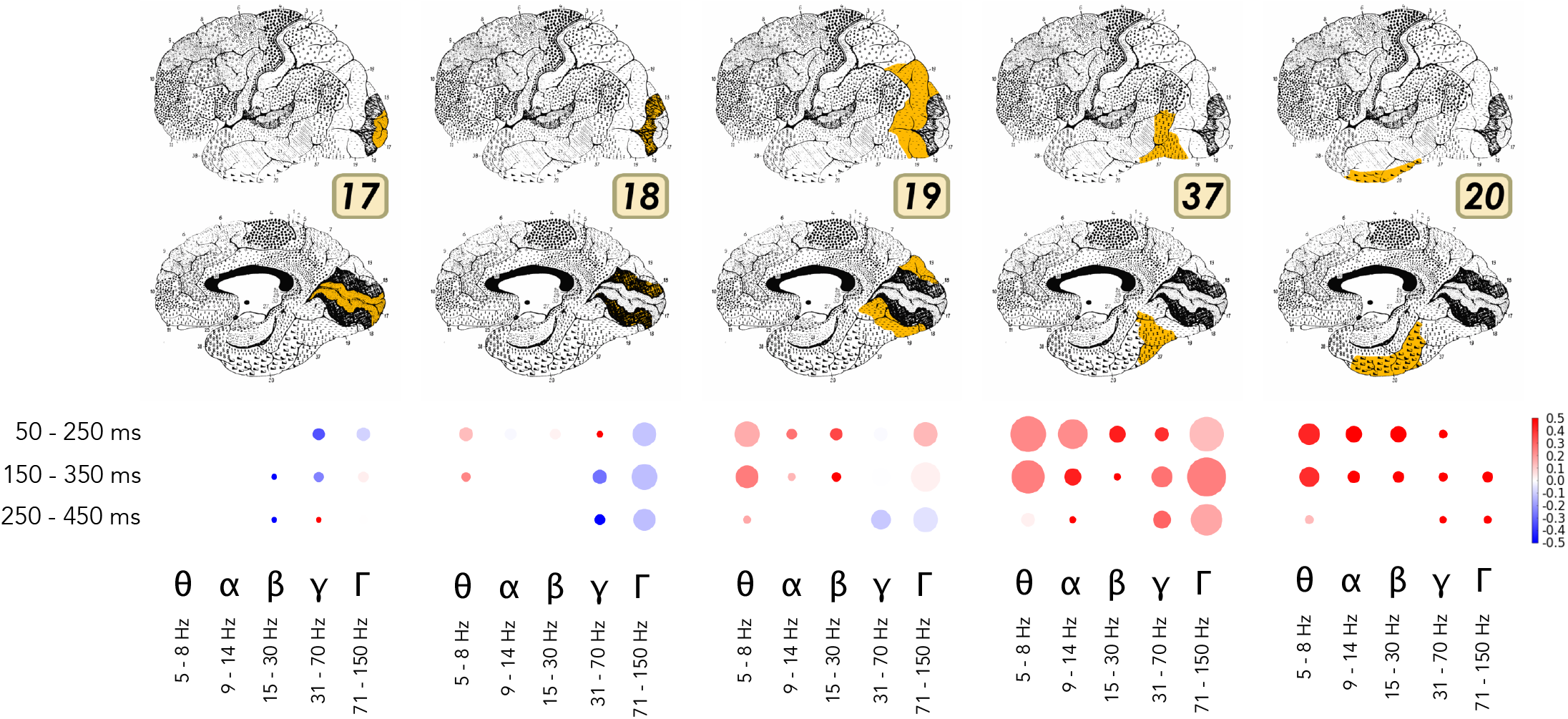
Area-specific analysis of volume of neural activity and complexity of visual features represented by that activity. Size of the marker shows the sum of correlation coefficients between the area and DCNN for each particular band and time window. Color codes the ratio of complex visual features to simple visual features, i.e. the comparison between the activity that correlates with the higher layers (`conv5, fc6, fc7`) of DCNN to the lower layers (`convl, conv2, conv3`). Intense red means that the activity was correlating more with the activity of higher layers of DCNN, while the intense blue indicates the dominance of correlation with the lower areas. If the color is close to white then the activations of both lower and higher layers of DCNN were correlating with the brain responses in approximately equal proportion.

### Gamma activity is more specific to convolutional layers, while the activity in lower frequency bands is more specific to fully connected layers

We analysed volume and specificity of brain activity that correlates with each layer of DCNN separately to see if any bands or time windows are specific to particular level of hierarchy of visual processing in DCNN. Figure 8 presents a visual summary of this analysis. In the “Methods” section we have defined total volume of visual activity in layers *L*. We used this measure to quantify the activity in low and high gamma bands. We noticed that while the fraction of gamma activity that is mapped to convolutional layers is high 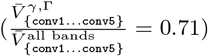, this fraction diminished in fully connected layers `fc6` and `fc7` 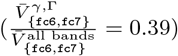. Note that `fc8` was excluded as it represents class label probabilities and does not carry information about visual features of the objects. On the other hand the activity in lower frequency bands (theta, alpha, beta) showed the opposite trend – fraction of volume in convolutional layers was 0.29, while in fully connected it grew to 0.61. This observation highlighted the fact that visual features extracted by convolutional filters of DCNN are more similar to gamma frequency activity, while the fully connected layers that do not directly correspond to intuitive visual features, carry information that has more in common with the activity in the lower frequency bands.

**Figure 8.**
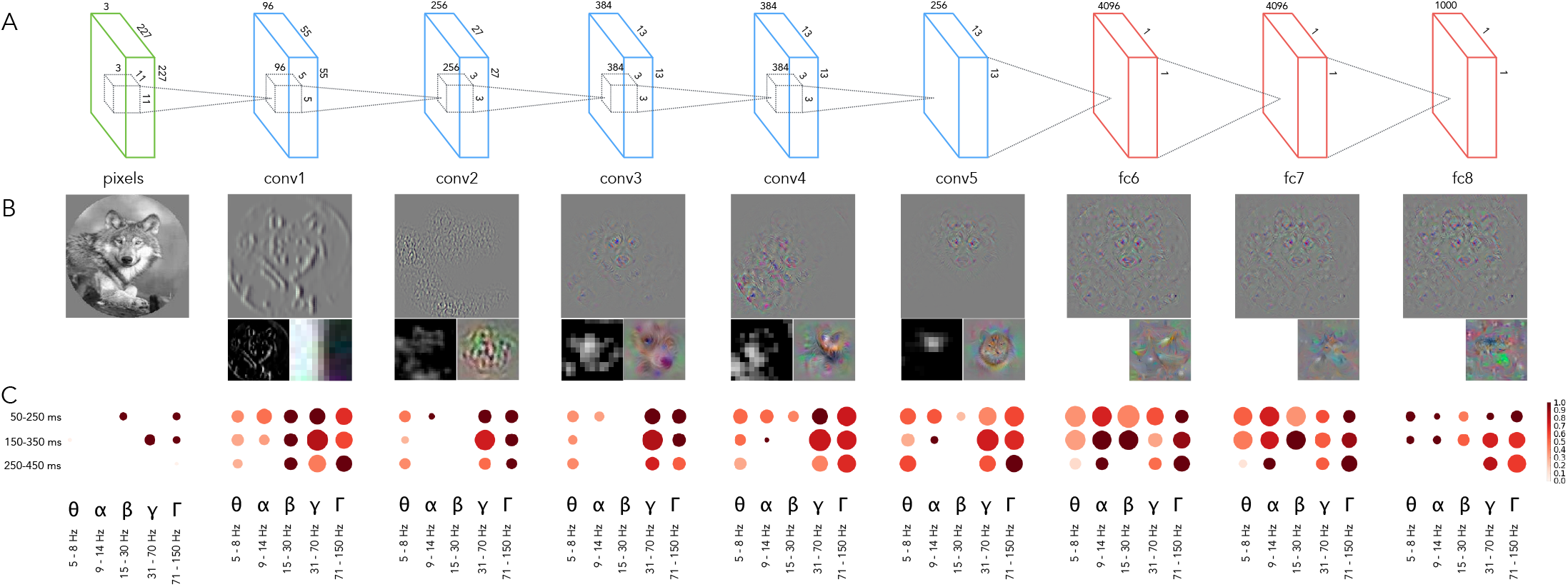
The architecture of the DCNN. Convolutional layer 1 consists of 96 feature detectors of size 11 × 11, they take as input pixels of the image and their activations create 96 features maps of size 55 × 55, architecture of all consecutive convolutional layers is analogous. Five convolutional layers are followed by 3 fully-connected layers of sizes 4096, 4096 and 1000 respectively. **B**. The leftmost image is an example input image. For each layer we have selected one interesting filter that depicts what is happening inside of the neural network and plotted: a) a reconstruction of the original image from the activity of that neuron using the deconvolution (Zeiler and Fergus, 2014) technique (upper larger image), (b) activations on the featuremap generated by that neuron (left sub-image) and (c) synthetic image that shows what input the neuron would be most responsive to (right sub-image). Visualizations were made with Deep Visualization Toolbox (Yosinski et al., 2015). All filters are canonical to AlexNet trained on ImageNet and can be explored using the above-mentioned visualization tool or visualized directly from the publicly available weights of the network. **C**. Specificity of neural responses across frequency bands and time windows for each layer of DCNN. Size of a marker is the total activity mapped to this layer and the intensity of the color is the specificity of the activity to the Brodmann areas constituting the ventral stream: BA17-18-19-37-20.

## Discussion

The recent advances in artificial intelligence research have been breathtaking. Not only do the deep neural networks match human performance in visual object recognition, they also provide the best model for how biological object recognition happens (Kriegeskorte, 2015; Yamins et al., 2013, 2014; Yamins and DiCarlo, 2016). Previous work has established a correspondence between hierarchy of the DCNN and the fMRI responses measured across the human visual areas (Güçlü and van Gerven, 2015; Eickenberg et al., 2016; Seibert et al., 2016; Cichy et al., 2016b). Further research has shown that the activity of the DCNN matches the biological neural hierarchy in time as well (Cichy et al., 2016b; Seeliger et al., 2017). Studying intracranial recordings allowed us to extend previous findings by assessing the alignment between the DCNN and cortical signals at different frequency bands. We observed that the lower layers of the DCNN explained gamma band signals from earlier visual areas, while higher layers of the DCNN, responsive for more complex features, matched with the gamma band signals from higher visual areas. This finding confirms previous work that has given a central role for gamma band activity in visual object recognition (Singer and Gray, 1995; Singer, 1999; Fisch et al., 2009) and feedforward communication (Van Kerkoerle et al., 2014; Bastos et al., 2015; Michalareas et al., 2016). Our work also demonstrates that the correlation between the DCNN and the biological counterpart is specific not only in space and time, but also in frequency.

### The role of gamma in object recognition

The research into gamma oscillations started with the idea that gamma band activity signals the emergence of coherent object representations (Gray and Singer, 1989; Singer and Gray, 1995; Singer, 1999). However, this view has evolved into the understanding that activity in the gamma frequencies reflects neural processes more generally. One particular view (Fries, 2005, 2015) suggests that gamma oscillations provide time windows for communication between different brain regions. Further research has shown that especially feedforward activity from lower to higher visual areas is carried by the gamma frequencies (Van Kerkoerle et al., 2014; Bastos et al., 2015; Michalareas et al., 2016). As the DCNN is a feedforward network our current findings support the idea that gamma rhythms provide a channel for feedforward communication. However, our results by no means imply that gamma rhythms are only used for feedforward visual object recognition. There might be various other roles for gamma rhythms (Buzsáki and Wang, 2012; Fries, 2015).

### Low vs high gamma in object recognition

We observed significant alignment to the DCNN in both low and high gamma bands. However, when directly contrasted the alignment was stronger for low gamma signals. Furthermore, for high gamma this alignment was more restricted in time, surviving correction only in the middle time window. Previous studies have shown that low and high gamma frequencies are functionally different: while low gamma is more related to classic narrow-band gamma oscillations, high frequencies seem to reflect local spiking activity rather than oscillations (Manning et al., 2009; Ray and Maunsell, 2011), the distinction between low and high gamma activity has also implications from cognitive processing perspective (Vidal et al., 2006; Wyart and Tallon-Baudry, 2008). In the current work we approached the data analysis from the machine learning point of view and remained agnostic with respect to the oscillatory nature of underlying signals. Importantly, we found that numerically the alignment to the DCNN was stronger and persisted for longer in low gamma frequencies. However, high gamma was more prominent when considering volume and specificity to visual areas. These results match well with the idea that whereas high gamma signals reflect local spiking activity, low gamma signals are better suited for adjusting communication between brain areas (Fries, 2005, 2015).

### Two groups of areas in the visual system are mapped to two groups of layers in the DCNN

We observed that the significant alignment depended on the fact that there are two groups of layers in the DCNN: the convolutional and fully connected layers. We found that these two types of layers have similar activity patterns (i.e. representational geometry) within the group but the patterns are less correlated between the groups (Figure 2). As evidenced in the data, in the lower visual areas (17,18) the gamma band activity patterns resembled those of convolutional layers whereas in the higher areas (37 and 20) gamma band activity patterns matched the activity of fully connected layers. Area 19 showed similarities to both types of DCNN layers.

Convolutional layers impose a certain structure on the networks connectivity – each layer consists of a number of visual feature detectors, each dedicated to finding a certain pattern on the source image. Each neuron of the subsequent layer in the convolutional part of the network indicates whether the feature detector associated with that neuron was able to find its specific visual pattern (neuron is highly activated) on the image or not (neuron is not activated). Fully connected layers on the other hand, as the name suggests, connect every neuron of a layer to every neuron in the subsequent layer, allowing for more flexibility in terms of connectedness between the neurons. The training process determines which connections remain and which ones die off. In simplified terms, convolutional layers can be thought of as feature detectors, whereas fully connected layers are more flexible: they do whatever needs to be done to satisfy the learning objective. It is tempting to draw parallels to the roles of lower and higher visual areas in the brain: whereas neurons in lower visual areas (17 and 18) have smaller receptive fields and code for simpler features, neurons in higher visual areas (like 37 and parts of area 20) have larger receptive fields and their activity explicitly represents objects (Grill-Spector and Malach, 2004; DiCarlo et al., 2012). On the other hand, while in neuroscience one makes the broad differences between lower and higher visual cortex (Grill-Spector and Malach, 2004) and sensory and association cortices (Zeki, 1993), this distinction is not so sharply defined as the one between convolutional and fully connected layers. Our hope is that the present work contributes to understanding the functional differences between lower and higher visual areas.

### Feedforward and feedback computations in object recognition

Visual object recognition in the brain involves both feedforward and feedback computations (DiCarlo et al., 2012; Kriegeskorte, 2015). What do our results reveal about the nature of feedforward and feedback compoments in visual object recognition? We observed that the DCNN corresponds to the biological processing hierarchy even in the latest analysed time-window (Figure 4). In a directly relevant previous work Cichy and colleagues compared DCNN representations to millisecond resolved MEG data from humans (Cichy et al., 2016b). There was a positive correlation between the layer number of the DCNN and the peak latency of the correlation time course between the respective DCNN layer and MEG signals. In other words, deeper layers of the DCNN predicted later brain signals. As evidenced on Figure 3 in (Cichy et al., 2016b), the correlation between DCNN and MEG activity peaked between ca 100 and 160 ms for all layers, but significant correlation persisted well beyond that time-window. In our work too the alignment in low gamma was strong and significant even in the latest time-window 250-450 ms, but it was significantly smaller than in the earliest time-window 50-250 ms. In particular, the alignment was the strongest for low gamma signals in the earliest time-window compared to all other frequency-and-time combinations.

### Limitations

The present work relies on data pooled over the recordings from 100 subjects. Hence, the correspondence we found between responses at different frequency bands and layers of DCNN is distributed over many subjects. While it is expected that single subjects show similar mappings (see also Figure 6), the variability in number and location of recording electrodes in individual subjects makes it difficult a full single-subject analysis with this type of data. We also note that the mapping between electrode locations and Brodmann areas is approximate and the exact mapping would require individual anatomical reconstructions and more refined atlases. Also, it is known that some spectral components are affected by the visual evoked potentials (VEPs). In the present experiment we could not disentangle the effect of VEPs from the other spectral responses as we only had one repetition per image. However, we consider the effect of VEPs to be of little concern for the present results as it is known that VEPs have a bigger effect on low frequency components, whereas our main results were in the low gamma band.

It must be also noted that the DCNN still explains only a part of the variability of the neural responses. Part of this unexplained variance could be noise (Güçlü and van Gerven, 2015; Khaligh-Razavi and Kriegeskorte, 2014). Previous works that have used RSA across brain regions have in general found the DCNNs to explain a similar proportion of variance as in our results (Cichy et al., 2016b; Seibert et al., 2016). It must be noted that the main contribution of DCNN has been that it can explain the gradually emerging complexity of visual responses along the ventral pathway, including the highest visual areas where the typical models (e.g. HMAX) were not so successful (Yamins et al., 2014; Khaligh-Razavi and Kriegeskorte, 2014). Recently it also has been demonstrated that the DCNN provides the best model for explaining responses to natural images also in the primate V1 (Cadena et al., 2017). Nevertheless, the DCNNs cannot be seen as the ultimate model explaining all biological visual processing (Kriegeskorte, 2015; Rajalingham et al., 2018). Most likely over the next years deep recurrent neural networks will surpass DCNNs in the ability to predict cortical responses (Kriegeskorte, 2015; Shi et al., 2017).

### Future work

Intracranial recordings are both precisely localized in space and time, thus allowing us to explore phenomena not observable with fMRI. In this work we investigated the correlation of DCNN activity with five broad frequency bands and three time windows. Our next steps will include the analysis of the activity on a more granular temporal and spectral scale. Replacing representation similarity analysis with a predictive model (such as regularized linear regression) will allow us to explore which visual features elicited the highest responses in the visual cortex. In this study we have investigated the alignment of visual areas with one of the most widely used DCNN architectures – AlexNet. The important step forward would be to compare the alignment with other networks trained on visual recognition task and investigate which architectures preserve the alignment and which do not. That would provide an insight into which functional properties of DCNN architecture are compatible with functional properties of human visual system.

### Conclusion

In the present work we studied which frequency components match the increasing complexity of representations of an artificial neural network. As expected by previous work in neuroscience, we observed that gamma frequencies, especially low gamma signals, are aligned with the layers of the DCNN. Previous research has shown that in terms of anatomical location the activity of DCNN maps best to the activity of visual cortex and this mapping follows the propagation of activity along the ventral stream in time. With this work we have confirmed these findings and have additionally established at which frequency ranges the activity of human visual cortex correlates the most with the activity of DCNN, providing the full picture of alignment between these two systems in spatial, temporal and spectral domains.

## Acknowledgements

We thank Martin Hebart and four reviewers for helpful comments. IK, RV and JA thank the financial support from the Estonian Research Council through the personal research grants PUT438 and PUT1476. This work was supported by the Estonian Centre of Excellence in IT (EXCITE), funded by the European Regional Development Fund.

